# Genomic prediction of cognitive traits in childhood and adolescence

**DOI:** 10.1101/418210

**Authors:** A.G. Allegrini, S. Selzam, K. Rimfeld, S. von Stumm, J.B. Pingault, R. Plomin

## Abstract

Recent advances in genomics are producing powerful DNA predictors of complex traits, especially cognitive abilities. Here, we leveraged summary statistics from the most recent genome-wide association studies of intelligence and educational attainment to build prediction models of general cognitive ability and educational achievement. To this end, we compared the performances of multi-trait genomic and polygenic scoring methods. In a representative UK sample of 7,026 children at age 12 and 16, we show that we can now predict up to 11 percent of the variance in intelligence and 16 percent in educational achievement. We also show that predictive power increases from age 12 to age 16 and that genomic predictions do not differ for girls and boys. Multivariate genomic methods were effective in boosting predictive power and, even though prediction accuracy varied across polygenic scores approaches, results were similar using different multivariate and polygenic score methods. Polygenic scores for educational attainment and intelligence are the most powerful predictors in the behavioural sciences and exceed predictions that can be made from parental phenotypes such as educational attainment and occupational status.

## Introduction

Ever increasing sample sizes and methodological advances in polygenic methods have made it possible to powerfully predict complex traits such as cognitive abilities without knowing anything about the causal chain between genes and behaviour. Progress in predicting cognitive traits from inherited DNA variants has been rapid in the past five years and especially in the past year ^1^. Three methodological advances have mainly been responsible for this progress: increasingly large genome-wide association (GWA) studies, genome-wide polygenic scores (GPS) and multivariate analytic tools. The key has been the recognition that the largest associations are extremely small, accounting for less than 0.05% of the variance ^2^. To achieve sufficient power to detect such small effect sizes, samples in the hundreds of thousands are needed before GWA studies can begin to detect these tiny effects. Because the largest associations are so small, useful predictions of individual differences can only be made by aggregating the effects of thousands of DNA variants in GPS ^3^. The third advance consists in the development of genomic methods that leverage genetic correlations between traits to boost power for variant discovery ^4^ and polygenic risk prediction ^5^.

Together, these three advances have greatly increased the ability to predict intelligence, educational attainment (years of schooling), and educational achievement (tested performance). For example, for intelligence, until 2017, no replicable associations were found in seven GWA studies ^6-12^, which we refer to collectively as ‘IQ1’. These studies had sample sizes from 18,000 to 54,000, which seemed large at the time but were not sufficiently powered to detect effect sizes of 0.05%. GPS derived from these IQ1 GWA studies at most accounted for 1% of the variance in independent samples. Increasing GWA sample sizes to 78,000 IQ2; ^13^ and then to 280,000 IQ3; ^14^ paid off in increasing predictive power of GPS from 1% to 3% to 4%. Here we present results for IQ3.

Educational attainment has led the way in terms of increasing GWA sample size, from 125,000 in 2013 (EA1^15^) to 294,000 in 2016 (EA2^16^) to 1.1 million in 2018 (EA3^17^). The growing sample sizes increased the predictive power of GPS from 2% to 3% to 12% of the variance in educational attainment ^1^. We showed that EA GPS predicted more variance in tested educational achievement than in the target trait of educational attainment. EA1 predicted 3% of the variance in educational achievement at age 16 ^18^ and EA2 predicted 9% of the variance for overall educational achievement at age 16 ^19a^ and up to 5% of the variance for reading ability at age 12 ^20b^.

Because ‘years of education’ is obtained as a demographic marker in most GWA studies, it was possible to accumulate samples sizes with the necessary power to detect very small effect sizes. It is more difficult to obtain very large sample sizes for intelligence, which needs to be assessed with a psychometric test administered to each individual, whereas years of education can be captured with a single self-reported item. Because of the large sample size available for EA GWA studies and the substantial genetic correlation between EA and intelligence, EA GPS predicted as much or more variance in intelligence than did GPS derived from GWA analyses of the target trait of intelligence itself. EA1 predicted 1% of the variance in intelligence ^18, 21^ and EA2 predicted 4% of the variance ^16^. Here we present results for EA3.

Finding that EA GPS predict educational achievement and intelligence better than do GWA of the target traits themselves suggests the usefulness of multivariate approaches. We used a multivariate GPS approach involving regularized regression to show that with EA2 and 80 other GPS we could predict 11% of the variance in educational achievement at age 16 and 5% of the variance in intelligence at age 12 ^22^. Although adding 1-2% to the predictive power of GPS might not seem like much, it should be noted that five years ago the total variance that could be predicted in either trait was 0%.

The aim of the present study is to estimate how much variance in intelligence and educational achievement can be predicted by applying several state-of-the-art multivariate genomic approaches and leveraging highly powered GWA summary statistics. First we compare three polygenic score methods (PRSice, LDpred, and Lassosum) and test how much variance the new IQ3 and EA3 GPS maximally predict. We then jointly analyse IQ3 and EA3 with three highly correlated traits to boost predictive power and compare performance of three multi-trait methods (Genomic SEM, MTAG and SMTpred) using predictive power as our criterion.

We conducted these analyses in a sample of 7,026 unrelated individuals from the Twins Early Development study, which is representative of the UK population ^23^. We analysed intelligence and educational achievement at the end of compulsory schooling in the UK at age 16, and at age 12 in order to investigate developmental trends in genomic prediction. Based on previous research ^19^, we expected genomic predictions to increase from 12 to 16.

## Materials and Methods

### Sample

The sample was drawn from the Twins Early Development Study (TEDS; ^24^, an ongoing population-based longitudinal study. It consists of twins born in England and Wales between 1994 and 1996, who have been assessed on a variety of psychological domains. More than 10,000 twin pairs representative of the general UK population ^23^ remain actively involved in the study to date. Ethical approval for TEDS has been provided by the King’ s College London ethics committee (reference: 05/Q0706/228). Parental consent was obtained before data collection. Genotypes for 10,346 individuals (including 3,320 DZ twin pairs) were processed using stringent quality control procedures followed by SNP imputation using the Haplotype Reference Consortium (release 1.1) reference panels. Current analyses were limited to the genotyped and imputed sample of 7,026 unrelated individuals. Following imputation, we excluded variants with minor allele frequency < 0.5%, Hardy-Weinberg equilibrium p-values of < 1×10^-5^. To ease computational demands, we selected variants with an info score of 1, resulting in 515,000 SNPs used for analysis (see Supplementary Methods S1 for a full description of quality control and imputation procedures).

### Outcome variables

The outcome variables were intelligence and educational achievement at ages 12 and 16. Intelligence was assessed as a composite of verbal and nonverbal web-based tests. Educational achievement was indexed by a mean of scores on the compulsory subjects of English, mathematics and science obtained from the UK National Pupil Database. A more detailed description of outcome variables is provided in the Supplementary Methods S2. Supplementary Table S1 includes descriptive statistics for the outcomes variables and Supplementary Figure S1 shows phenotypic correlations. Phenotypes and polygenic scores were corrected for age, sex and 10 genetic principal components. The obtained standardised residuals were used in all subsequent analyses.

### Discovery sets - GWA summary statistics

We based our prediction models on beta weights derived from large, publicly available, GWA summary statistics. Of central importance for our analyses were the most recent GWA studies of educational attainment EA3 ^17^ and intelligence IQ3 ^14^. The original IQ GWA meta-analysis included TEDS as one of its samples, therefore, to avoid bias due to sample overlap with our target sample, we used summary statistics from new GWA analyses that excluded TEDS. The EA3 summary statistics employed here do not include 23andMe data (∼300k individuals) due to their data availability policy.

### Polygenic score approaches

We used IQ3 and EA3 summary statistics to construct genome-wide polygenic scores (GPS) comparing three distinct approaches: PRSice2 ^25^, a clumping/pruning + P-value thresholding (P+T) approach, with an in-built high-resolution option that returns the best-fit GPS for the trait of interest; LDpred ^26^, a Bayesian approach that uses a prior on the expected polygenicity of a trait (assumed fraction of non-zero effect markers) and adjusts for linkage disequilibrium based on a reference panel to compute SNPs weights; and Lassosum ^27^, a machine-learning approach which uses penalized regression on GWA summary statistics to produce more accurate beta weights.

A detailed description of the construction of these polygenic scores is included in Supplementary Methods S3.

### Multivariate approaches

In order to boost power of IQ3 (N = 266,450) and EA3 (N = 766,345) GWA results and thus precision of beta weights to construct more predictive IQ3 and EA3 polygenic scores, we jointly analysed these summary statistics with three cognitive and educationally relevant traits: “Income” ^28^ (N = 96,900), “Age when completed full time education” ^29^ (N = 226,899) and “Time spent using computer” ^29^ (N = 261,987). The choice of these traits is consistent with a multi-trait framework, as these traits show the highest genetic correlations with IQ and educational attainment among publicly available GWA summary statistics, with pairwise-genetic correlations ranging from ∼.5 to ∼.9 (see Supplementary Figure S2). Summary statistics from these GWA studies are reported in Supplementary Table S2.

We used three recently developed multi-trait methods, one of which is specifically designed to boost polygenic score prediction: SMTpred ^5^, and two of which are strictly speaking multivariate GWA approaches, designed to boost power for discovery, but which have been shown to boost predictive power of polygenic scores created from multi-trait reweighted summary statistics: MTAG ^4^ and Genomic SEM ^30^. Details about these methods are provided in Supplementary Methods S4. Briefly, SMTpred ^5^ is a multi-trait extension of the random effects model approach, which can be used to create multivariate best linear unbiased predictors based on summary statistics (wMT-SBLUP). MTAG is a generalization of inverse-variance weighted meta-analysis, which jointly analyses univariate GWA summary statistics. It boosts power for discovery for each trait conditional on the other traits and outputs trait-specific summary statistics. Genomic SEM is a two-stage structural equation modelling approach that can be applied in the context of multivariate GWA. In the form employed here (common factor GWA analysis) it directly tests effect of SNPs on a latent genetic factor defined by several indicators (i.e. traits) and outputs summary statistics for the common factor. We also compared these new multivariate approaches to a simple multiple regression on intelligence and on educational achievement using five GPS, each derived from the univariate GWA summary statistics used in multi-trait analyses.

## Analyses

### Univariate analyses

We first calculated polygenic scores for the IQ3 and EA3 GWA summary statistics using PRSice, LDpred and Lassosum. This was done to compare current state-of-the-art polygenic scores approaches and in order to obtain a benchmark against which to compare improvements in prediction accuracy due to multivariate GWA analyses. For each phenotype (i.e. intelligence and educational achievement at ages 12 and 16), we randomly split the sample into training and test sets (∼50% training, ∼50% test). Supplementary Table S1 shows descriptive statistics for each set. In the training sets, parameter optimization of GPS was performed, in which each GPS instrument (or p-value threshold in the case of PRSice, fraction of markers with nonzero effect in the case of LDpred, and tuning parameters in the case of Lassosum) was tested on each of the four phenotypes and the best instrument was selected with respect to prediction accuracy (as indexed by R^2^). Performance of the optimized GPS instrument retained from the validation was then assessed in the test sample in order to evaluate how well the chosen predictors would perform in independent samples. We then proceeded to perform the multi-trait analyses.

### Multivariate analyses

We performed a multi-trait reweighting in SMTpred after transforming the ordinary least square betas from GWA studies of ‘IQ’, ‘EA’, ‘Income’, ‘Age when completed full time education’ and ‘Time spent using computer’ in approximate Best Linear Unbiased Predictors (BLUP) using GCTA-Cojo **^31^.** We then used LDSC to calculate SNP h2 and genetic correlations between traits and proceeded to the multivariate weighting of traits as described in (Meier et al., 2018) to obtain multi-trait summary statistics BLUP (wMT-SBLUP; see also Supplementary Methods S4).

MTAG was run on the five GWA summary statistics (IQ, EA, Income, Age completed full time Education, Income) using standard settings. Because MTAG combines differently powered summary statistics (as indexed by the GWAS mean χ^2^; see Supplementary Methods S4), as well as differing degrees of genetic overlap between traits, it can lead to an increased rate of false positives Type I error ^4^. However, this is not an issue in the present study, which focuses on prediction accuracy rather than variant discovery. It has been shown ^4^ that MTAG estimates consistently have a lower genome-wide mean-squared error compared to single-trait GWA estimates, and, therefore, polygenic scores created from MTAG perform better than those created at the univariate level. However, in order to control for type I error inflation, we used the recommended ^4^ false discovery rate (FDR) calculations (see Supplementary Methods S4).

The same five summary statistics were analysed using Genomic SEM. First a common factor model with the five summary statistics as indicators was fitted using a weighted least-square (WLS) estimator (default setting in Genomic SEM). Then a common factor GWA analysis with a WLS estimator was run, testing effects of single SNPs on the common factor. The WLS estimator was expected to yield lower standard errors and possibly increased prediction accuracy of GPS ^30^.

We then created polygenic scores from the MTAG EA, MTAG IQ and common factor GWA summary statistics across the three polygenic scores approaches, after splitting the sample into a training set to tune parameters and a testing set to assess model performance. In the case of SMTpred, polygenic scores for IQ3 and EA3 converted and reweighted indices (wMT-SBLUP) were calculated using PLINK ^32^. These multi-trait predictors were then directly tested for model performance in the test set, as with the other GPS approaches.

## Results

### Polygenic score prediction of IQ and EA across GPS methods

Figure 1 shows variance in intelligence and educational achievement predicted by IQ3 GPS and EA3 GPS calculated following three polygenic score methods (PRSice, LDpred and Lassosum). Supplementary Table S3 presents associations in the training and test sets across all models.

**Figure 1.**
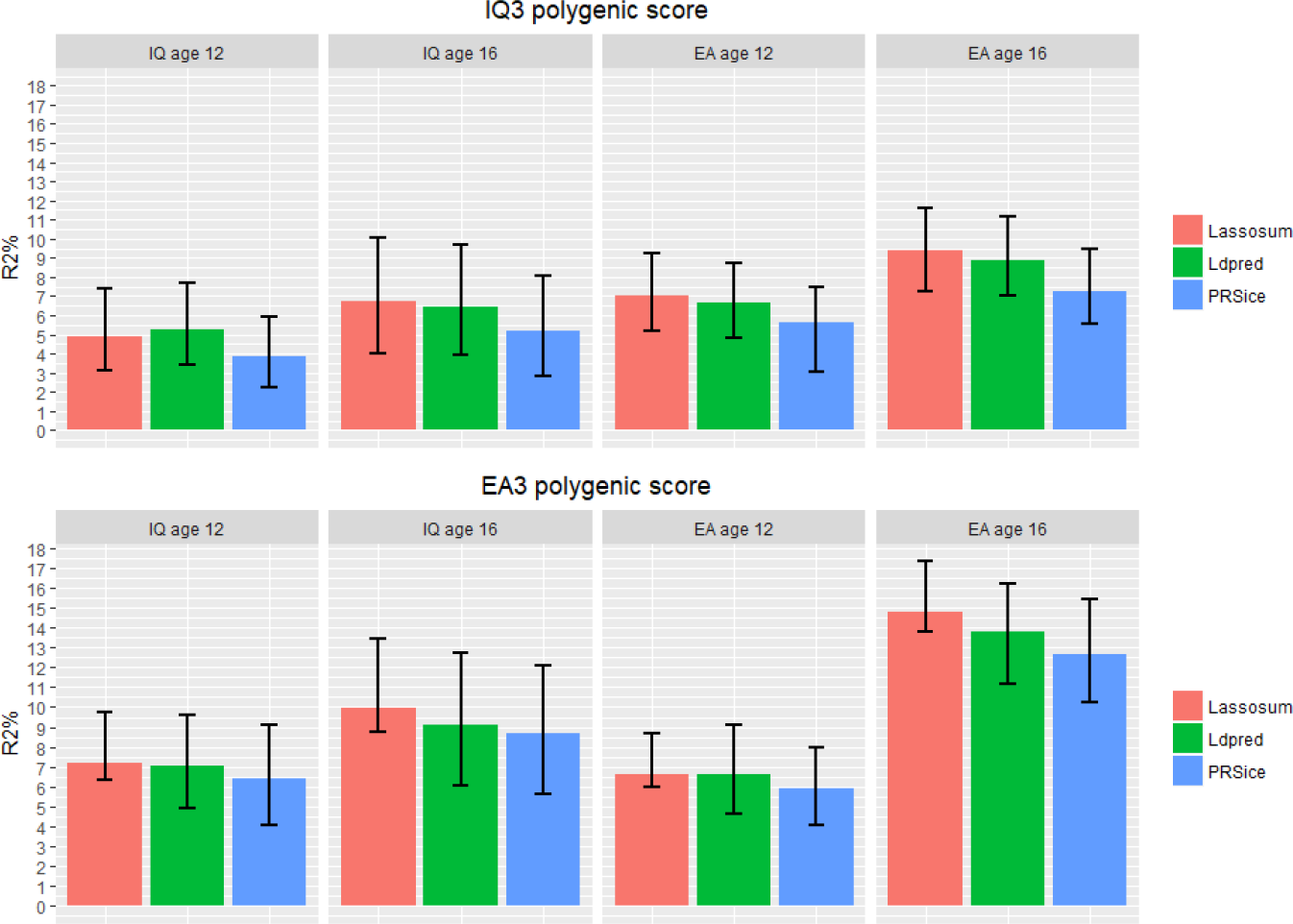
Polygenic score prediction of intelligence (IQ) and educational achievement (EA) at age 12 and 16. Figure shows polygenic prediction accuracy across polygenic score methods. Error bars are bootstrapped 95% confidence intervals based on 1,000 replications.

For intelligence, IQ3 GPS predicted a maximum of 5.3% (β = 0.221, se = 0.023, p < .0001) of the variance at age 12 and 6.7% (β = 0.266, se = 0.032, p < .0001) at age 16. For educational achievement, EA3 GPS predicted a maximum of 6.6% (β = 0.259, se = 0.020, p < .0001) of the variance at age 12 and 14.8% (β =0.389, se = 0.019, p < .0001) at age 16. EA3 GPS was also a powerful predictor of intelligence, predicting 7.2% (β = 0.265, se = 0.024, p < .0001) of the variance in intelligence at age 12 and 9.9% (β = 0.321, se = 0.031, p < .0001) at age 16.

Generally, Lassosum was the most powerful approach, predicting up to 1% more of the variance compared to LDpred and up to 2% more compared to PRSice. It is of note, however, that bootstrapped 95% confidence intervals consistently overlapped across prediction estimates.

### Multi-trait polygenic score prediction

Results of multivariate GWA analyses are reported in Supplementary Methods S4 and supplementary Tables S5 and S6. Here we report results of polygenic score associations for our best predictive polygenic models after multi-trait approaches were applied to GWA summary statistics. Supplementary Figure S4 shows a comparison of variance predicted in intelligence and educational achievement at ages 12 and 16 in the test samples across polygenic score methods after multi-trait analyses. Supplementary Table S4 reports details of these results.

Figure 2 shows variance predicted in intelligence and educational achievement at age 16 by polygenic scores derived from multi-trait methods. For intelligence, variance predicted by IQ3 GPS increased from 6.7% (Figure 1) to a maximum of 10.0% (β = 0.327, se = 0.032, p < .0001) at age 16. For educational achievement, variance predicted by EA3 GPS increased from 14.8% to a maximum of 15.9% (β = 0.403, se = 0.018, p < .0001) at age 16. Again, EA3 GPS was generally the best performing predictor across phenotypes, predicting a maximum of 10.6% (β = 0.332, se =0.031, p < .0001) in intelligence. Similar improvements in prediction were observed at age 12 (see Supplementary Table S4 and supplementary figure S4).

**Figure 2.**
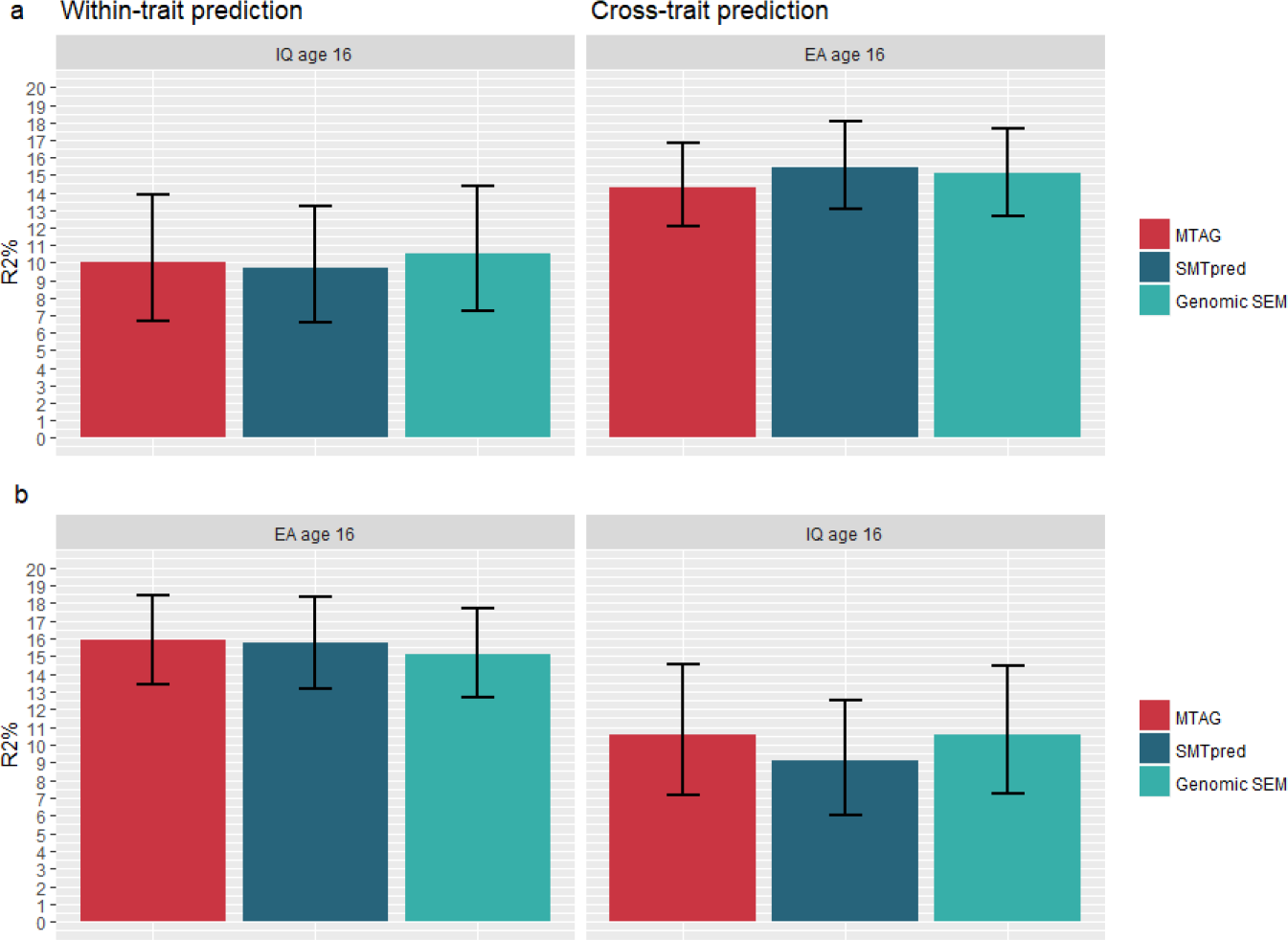
Within-trait and cross-trait polygenic score prediction of intelligence and educational achievement at age 16 across multi-trait methods. Note. MTAG = MTAG IQ3 (panel **a**)/ MTAG EA3 (panel **b**) polygenic scores constructed in Lassosum; SMTpred = IQ3 (panel **a**)/EA3 (panel **b**) wMT-SBLUP predictors; Genomic SEM = Common Factor polygenic score constructed in Lassosum (panel **a** and **b**). Error bars are bootstrapped 95% confidence intervals based on 1,000 replications.

### Polygenic scores quantile differences

Figure 3 shows the results for the best predictive models at age 16 by GPS deciles. For both intelligence (panel a) and educational achievement (panel b), the relationship with GPS deciles is linear and the lowest and highest deciles differ substantially. For intelligence, the mean difference (∼1 SD) is comparable to 15 IQ points. For educational achievement, the mean difference corresponds to an average ‘C’ grade for the lowest decile and an average ‘A’ grade for the highest decile. However, individuals in the lowest and highest deciles overlap considerably, as would be expected from GPS correlations of ∼0.32 with intelligence and ∼0.40 with educational achievement.

**Figure 3.**
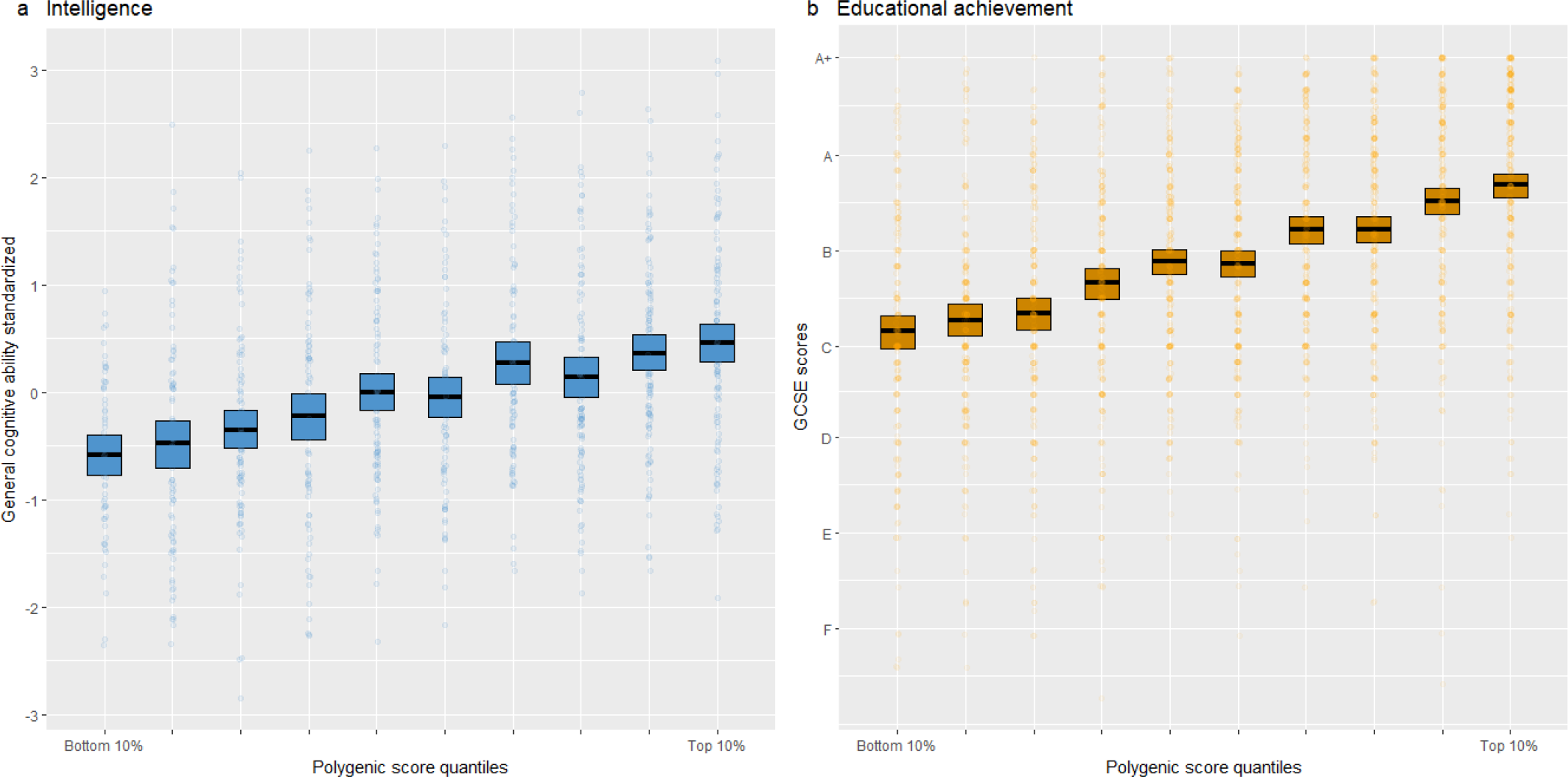
Mean intelligence scores (panel **a**) and mean educational achievement (panel **b**; GCSE grades) at age 16 by GPS deciles for the best polygenic predictors in the test set. Bars represent bootstrapped 95% confidence estimates.

### Sex differences

We tested associations for the best predictive model (i.e. MTAG EA3 GPS calculated in Lassosum) separately for males and females in the test set. For intelligence at age 16, the GPS predicted 10.73% of the variance (95% CI = 6.33 – 16.74) in males (N= 369, β = 0.334, se = 0.049) and 10.51% (95% CI = 6.49 – 15.41) in females (N = 558, β = 0.329, se = 0.040). For educational achievement in males (N = 1,105) the GPS predicted 14.2% (95% CI 10.96 – 17.86) of the variance (β = 0.375, se = 0.027); in females (N = 1,300) estimates were 17.24% (95% CI 13.51 – 21.43; β = 0.420, se = 0.025). To test the significance of these sex differences, we performed a Fisher’s r to z transformation of corresponding correlation coefficients. Sex differences were not significant for neither intelligence (observed z = 0.029, p = 0.488) nor educational achievement (observed z = −1,127, p= 0.129).

### Multiple regression model

We compared the results from our multi-trait GPS analyses to a simple multiple regression analysis using the five GPS from summary statistics of our multi-trait analyses (IQ, EA, income, age when completed full time education, time spent using computer) to predict intelligence and educational achievement. The multiple regression model predicted similar amounts of variance as the best single multi-trait GPS predictors. For intelligence, the adjusted R^2^ was 8.6% at age 12 and 9.9% at age 16. For educational achievement, the adjusted R^2^ was 9.6% at age 12 and 16.7% at age 16. Results are shown in Supplementary Table S7.

## Discussion

Using summary statistics from the latest GWA studies of intelligence (IQ3) and educational attainment (EA3), we report the strongest polygenic prediction estimates for cognitive-related traits to date. Comparing standard polygenic score approaches, we showed that IQ3 GPS predicts a maximum of 6.73% of the variance in intelligence at age 16, while EA3 GPS predicts 14.78% % of the variance in educational achievement at age 16.

In an attempt to boost predictive power, we compared results using state-of-the-art genomics methods that leverage the multivariate nature of traits in order to increase power of GWA summary statistics. We then tested boosted summary statistics across a number of polygenic score approaches, showing that we can predict 10.6% of the variance in intelligence and 15.9% of the variance in educational achievement, both at age 16. These results compare favourably with polygenic prediction estimates from the recent EA3 GWA analysis, whereby a polygenic score constructed from multi-trait summary statistics of educational attainment and three cognitive-related phenotypes predicted up to 13% of the variance in educational attainment and up to 10% in cognitive performance ^17^, this is especially notable given the larger discovery sample size employed in that study (N ∼ 1.1 million including 23andMe).

We found that trait prediction increased from age 12 to age 16. Polygenic scores become more predictive with age, probably because as the sample approaches adulthood it is closer in age to the samples in which beta weights were estimated in the original GWA studies for IQ3 and EA3. Another possible reason for this finding is that given that heritability of intelligence increases with age ^33^, also does the variance that can be predicted by cognitive related polygenic scores. Lastly, we did not find significant differences in the predictive power of IQ3 and EA3 for males and females.

These results indicate the usefulness of taking into account the multivariate nature of complex traits in polygenic prediction, and add to the possibility of practical use of polygenic scores at the level of individuals ^34^. It is important to note that we randomly split our sample (∼50%) to validate our models and assessed performance of prediction models in the test sample in order to avoid overfitting. Because TEDS is a fairly representative sample of the UK population, these prediction estimates are expected to be a close representation of how these models would perform in other, similar, samples. Overall, multi-trait methods were successful in increasing variance predicted, compared to our ‘baseline’ predictions estimates increased on the order of 1% to 3%. Multi-trait methods were especially useful in increasing predictive power of the IQ3 GPS, which was constructed using a less powerful summary statistic than the EA3 GPS. However, differences in prediction accuracy across the tested combinations of genomic methods seemed to reflect differences in polygenic score approaches rather than in multi-traits approaches. Yet, reassuringly, there were no dramatic differences in prediction accuracy across polygenic score approaches either, especially when considering only approaches that do not perform clumping (thereby losing information across the genome).

One limitation that could affect the interpretability of our findings is that MTAG, by jointly analysing traits with differing levels of power and genetic overlap, might confound the genetic architecture of boosted traits with that of other traits. In this regard we performed the recommended False discovery rate corrections which suggested that this was not an issue for IQ3 (FDR < .001) or EA3 (FDR <.0001). Finally, a general limitation of all genomic analyses is that they only assess additive effects of common SNPs used on currently SNP arrays. SNP heritability is the ceiling for polygenic score prediction, which is about 20%^14^ of the total variance for intelligence and 30% ^35^ for educational achievement. Viewed in this light, our best polygenic scores predict about half of the SNP heritability. With bigger and better GWA studies and other methodological advances like multivariate approaches, the missing SNP heritability gap will be narrowed. Polygenic scores will only reach their full potential when we are able to close the gap between SNP heritability (about 25%) and family study estimates of heritability (about 50%).

Nonetheless, these polygenic scores predictions are already among the strongest predictors in the behavioral sciences. For example, in TEDS, these polygenic scores predict educational achievement better than does parental educational attainment (14% for mothers, 15% for fathers) or occupational status (5% for mothers, 6% for fathers). Because inherited DNA variants do not change during development, polygenic scores are unique predictors in two ways. First, unlike other characteristics of the individual, DNA variants can predict individual differences in adult behaviour from birth. Second, unlike other correlations, associations between DNA variants and behaviour are causal from DNA to behaviour in the sense that there can be no backward causation from behaviour to DNA. These unique features will put genomic prediction of cognitive traits in the front line of the DNA revolution.

## Acknowledgements

We gratefully acknowledge the ongoing contribution of the participants in the Twins Early Development Study (TEDS) and their families. TEDS is supported by a program grant to RP from the UK Medical Research Council (MR/M021475/1 and previously G0901245), with additional support from the US National Institutes of Health (AG046938). The research leading to these results has also received funding from the European Research Council under the European Union’ s Seventh Framework Programme (FP7/2007-2013)/ grant agreement n^°^ 602768 and ERC grant agreement n^°^ 295366. RP is supported by a Medical Research Council Professorship award (G19/2).

## Author contributions

AGA and RP conceived and designed the study. AGA analysed and interpreted the data. AGA and RP wrote the manuscript. SS, KR, SvS and JBP contributed to and critically reviewed the manuscript.

